# Low Carbohydrate and Low Fat Diets with Equal Protein Content Lead to Similar Improvements in Body Composition and Glucose Tolerance in Obese Mice subjected to Caloric Restriction

**DOI:** 10.1101/830752

**Authors:** Petras Minderis, Andrej Fokin, Mantas Dirmontas, Aivaras Ratkevicius

## Abstract

**Background:** Reported differences in effects of low and high carbohydrate diets on weight control and metabolic health are controversial. We aimed to examine if such diets induce different improvements in body composition and glucose tolerance under conditions of caloric restriction (CR) in obese mice.

**Methods:** Male C57BL/6J mice (n = 20) were fed obesogenic diet (45 and 17.5% kcal from fat and sugar) *ad libitum* for 18 weeks and then subjected to 6-week CR which progressively increased up to 40% using either Low Fat diet (20, 60, 20% kcal from fat, carbohydrate, protein, n = 10) or Low Carb diet (20, 60, 20% kcal from carbohydrate, fat, protein, n = 10). Mice fed regular chow diet *ad libitum* served as controls (n = 10). Body mass, hind limb muscle mass, fat mass, energy expenditure and glucose tolerance were compared between the groups.

**Results:** Low Fat and Low Carb groups had similar body mass (*p* > 0.05) prior to CR which was 30% greater compared to control group (*p* < 0.001). CR resulted in weight loss with no differences between Low Fat and Low Carb groups (30.0 ± 5.6 and 23.8 ± 7.5%, *p* > 0.05). Weight loss was mainly due to fat loss in both groups. Energy expenditure of freely moving mice did not differ between the groups (*p* > 0.05). Intraperitoneal glucose tolerance test improved compared to control group (*p* < 0.05) and values before CR (*p* < 0.01) but without differences between Low Fat and Low Carb groups (*p* > 0.05).

**Conclusions:** Dietary carbohydrate or fat content when protein is equated does not play a significant role for body composition and metabolic health benefits of caloric restriction in obese mice.

## Introduction

Obesity is a risk factor for many non-infectious chronic diseases including cardiovascular heart disease, stroke, diabetes, and cancer which are the major causes of premature death in many countries around the world (1–4). Prevalence of obesity is steadily increasing (5) and becoming a threat to economic prosperity and national security as identification of solutions to obesity epidemic is high on the agenda worldwide (6, 7).

According to the paradigm of energy balance animals and humans gain weight when their energy intake exceeds energy expenditure (8). Increase in physical activity could prevent weight gain, but adjustments in diet are often easier to implement on the population level (9). A key question is what diet is best suited for weight control. A popular believe is that macronutrient composition of food are also important alongside prerequisite caloric restriction (10). Indeed, effect on satiety and dietary-induced thermogenesis are greater for dietary protein compared to carbohydrates or fat (11, 12). Human overfeeding studies suggest that protein has a smaller detrimental effect on body composition compared to carbohydrates and fat which are usually the major candidates for restriction in various diets for weight control (13). It is still controversial whether proportions of these two macronutrients are important for metabolic health. One of the theories proposes that dietary carbohydrates are inherently more obesogenic than fat due to strong effect on insulin secretion (14). The so-called carbohydrate-insulin model of obesity is criticized as lacking strong evidence in support of it (15). Nevertheless, a recent randomized-controlled study with humans demonstrated that energy expenditure was by up to 478 kcal per day greater on a low carbohydrate diet compared to high carbohydrate diet for a similar energy intake (16). Thus, diets stimulating energy expenditure while keeping unchanged energy input side would be a promising strategy in successful weight management. However, concerns were raised about suitability of doubly labelled water technique to measure energy expenditure in diets of varying carbohydrate and fat distribution as in above mentioned study of Ebbeling et al. (2018) (16–18). Human nutritional epidemiologic research addressing comparisons of different composition diets have also been plague by methodological difficulties which mainly concern assessment of food intake (19).

It appears that inbred mouse model is well suited to examine the controversial issue about the importance of dietary composition for health outcomes. Key advantage of such studies is that food intake can be controlled much better than in human studies and unpredictable effects of genetic factors are minimized. C57BL/6J mouse strain is prone to obesity when fed *ad libitum* (20) and tolerate well various diets with large differences in carbohydrate and fat content (21, 22). A recent study of 29 diets has demonstrated that dietary fat content was associated with greater energy intake and preponderance to obesity in these mice fed *ad libitum* (23). Our aim was to compare changes in body composition and metabolic adaptations of C57BL/6J mouse strain in response to two energy-restricted diets with large differences in carbohydrate and fat content (24). We hypothesize that changes in body composition would not differ between these two diets if they match for the total caloric content and protein-derived calories.

## Materials and methods

### Animals and experiments

The study was carried out at the Lithuanian Sports University with approval of all the procedures by the Lithuanian State Food and Veterinary Service in 2018 (Ref. # G2 – 90). The breeding pairs of C57BL/6J mouse strain were obtained from the Jackson laboratory (Bar Harbor, Maine, USA) and male mice were used in the experiment. Mice were housed at ambient temperature 20–21 °C and 40–60% humidity with an alternating 12-h light/dark cycle. After the weaning mice were housed two to five animals per cage and fed *ad libitum* with a regular grain-based rodent chow diet (Kombi, Joniskio grudai, Lithuania) and had unrestricted access to a tap water. At 10 weeks of age mice (n = 30) were switched to obesogenic high fat and sugar diet (D12451, 45 and 17.5% kcal from fat and sugar, Research Diets, New Brunswick, NJ, USA) for 18 weeks (25). This was followed by 6-week caloric restriction (CR) on either low fat diet (Low Fat, n = 10) or low carbohydrate diet (Low Carb, n = 10). Ten weight-matched mice prior CR were examined as pre-diet obese controls (Pre).

### Dietary intervention

#### Obesity phase

After 10 weeks of 18-week exposure to the obesogenic diet mice were moved into separate cages and food consumption was assessed every week for each mouse by subtracting food leftovers from initially provided food with corrections for humidity effect on the pellets weight. Daily energy intake (DEI) of mice was calculated as follows:

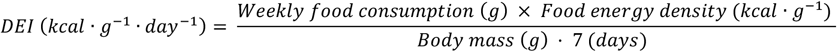

Three-week average of DEI of our mouse colony was 0.42 ± 0.4 kcal · g^−1^ · day^−1^ and only mice gaining at least 20% of weight compared to the age-matched group on the regular chow diet (Regular, n = 10) were used for CR study.

#### Calorie restriction (CR) phase

28-week-old obese mice were randomly assigned to one of the two CR groups and Pre group which was used for assessment of body composition and metabolism at the baseline. During 6 weeks CR was gradually increased from 20% (1 week) to 30% (2-4 week) and 40% (5-6 week) of the calculated energy intake on the *ad libitum* obesogenic diet. Energy intake during CR phase was estimated for each mouse individually by reducing DEI by the extent of caloric deficit and multiply it by initial body mass of the animal prior CR. The amount of food was corrected for different caloric density of the diets (4.1 and 5.2 kcal · g^−1^ for Low Fat and Low Carb, respectively) to achieve equal total energy and protein content in the diets, i.e. 20, 60 and 20% kcal from fat, carbohydrate and protein for Low Fat (D17100401, Research Diets, New Brunswick, NJ, USA) and 20, 60 and 20% kcal from carbohydrate, fat and protein for Low Carb (D12492, Research Diets, New Brunswick, NJ, USA), respectively. Details of the macronutrient composition and sources of the diets are presented in Supplementary table 1.

### Glucose tolerance

A 6-time point intraperitoneal glucose tolerance tests (IPGTT) were carried out after an overnight fasting during the final 6th week of CR. IPGTT began at 8:00 – 9:00 a.m. Mice were subjected to an intraperitoneal injection of glucose solution (2 g glucose · kg body wt^−1^) and glucometer (Glucocard X-mini plus GT-1960, Arkray, Japan) was used to measure glucose in the whole blood samples from the tail vein at 0, 15, 30, 60, 90 and 120 min after injection. The area under curve (AUC) of glucose response in IPGTT was calculated using Prism 6.0 software (GraphPad Software Inc., CA, USA).

### Body composition

During CR mice were weighed weekly with a precision of 0.1 g (440-45N, Kern, Germany). At the end of this phase, mice being at 34 weeks of age were euthanized with an inhalation of CO_2_. Immediately following this procedure skeletal muscles and body fat were sampled and weighed with a precision of 0.1 mg (ABS 80-4, Kern, Germany). Combined hindlimb muscle mass was calculated as a sum of the gastrocnemius, plantaris, soleus, tibialis anterior and extensor digitorum longus muscle mass. The muscles were trimmed from all visible tendons and blotted dry just before weighing. Combined body fat mass was assessed as the sum of the hindlimb white adipose subcutaneous (sWAT), gonadal (gWAT), mesenteric (mWAT), perirenal (pWAT) and intrascapular brown adipose tissue (iBAT) as in previous studies (26, 27).

### Energy expenditure and physical activity

Mice were fasted overnight and indirect calorimetry for assessment of total energy expenditure was applied together with measurements of physical activity in freely moving mice during the final week of CR using methods described in our previous study (28). Briefly, the metabolic cage of standard size was connected to the gas analyser (LE405, Panlab Harvard Apparatus, Spain) and the switching device (LE400, Panlab Harvard Apparatus, Spain) for the control of the air flow. Gas analyser was calibrated at the high point (50% O_2_, 1.5% CO_2_) and at the low point (20% O_2_, 0% CO_2_). Air flow was set to 250 ml · min^−1^ with 3-min switching time between measurements of O_2_ and CO_2_ concentrations in the metabolic cage and the external environment. All metabolism measurements were performed during a light period (from 9:00 a.m. to 3:00 p.m.). Each mouse was weighed (ABS 80-4, Kern, Germany) and transferred into metabolic cage for 3-h measurements with no food and water provided. Afterwards the mouse was weighed again and transferred back to the home cage with food and water supply. Total energy expenditure and respiratory quotient were calculated as the average values of the last 2-h of measurements (Metabolism software version 1.2, Panlab Harvard Apparatus, Spain) which is based on standard methods (29). Physical activity of mice was assessed using strain gauges mounted on the supporting constructions of the metabolic cage. The integral of ground reaction forces was used as an indirect measure of physical activity. The rearing was also assessed as lifts of the mouse body above infrared barriers set at 10 cm height.

### Statistical analysis

All data are presented as means ± SD or means with plotted individual data points. The statistical analysis was performed using Prism 6.0 and IBM SPSS Statistics v20 software. Normality of data distribution was verified with Shapiro-Wilk test. Means were compared with one-way analysis of variance (ANOVA) using Bonferroni’s *post hoc* test to assess differences between the studied groups of mice. Non-parametric Kruskal–Wallis test with Dunn’s *post hoc* analysis was applied in the cases when means did not meet a criterion of normal distribution. Two-way repeated measures ANOVA was used for analysis of body mass change when it was assessed repeatedly on the same animals. Analysis of covariance (ANCOVA) was applied using linear models to assess effects of mouse groups on energy expenditure as previously recommended for this type of analysis (30). In this case body mass and physical activity were used as covariates. Linear regression analysis was also used on the plots of energy expenditure over physical activity. Pearson’s correlation coefficient was calculated to assess strength of the association between the variables. The level of significance was set at *p* < 0.05.

## Results

### Energy intake was similar in Low Fat and Low Carb groups

Data on energy intake is presented in Fig. 1. We aimed at maintaining similar energy intake in Low Fat and Low Carb groups during CR. However, Low Carb group did not consume all the food during the first week of CR, and the unconsumed food was left in the feeder with subsequent daily portion added on top of the leftovers. However, after two weeks of CR, Low Carb group matched Low Fat group for energy intake. For the entire 6-week CR, these groups did not differ in the absolute (Fig. 1a) or body mass normalized energy intake when body mass before the start of CR was used for normalization (12.0 ± 0.4 and 12.1 ± 0.3 kcal · g initial body wt^−1^ for Low Fat and Low Carb, respectively; *p* > 0.05) (Fig. 1b). Mice in the Regular diet group had ~30% greater (*p* < 0.01) energy intake during the same period.

**Fig. 1.**
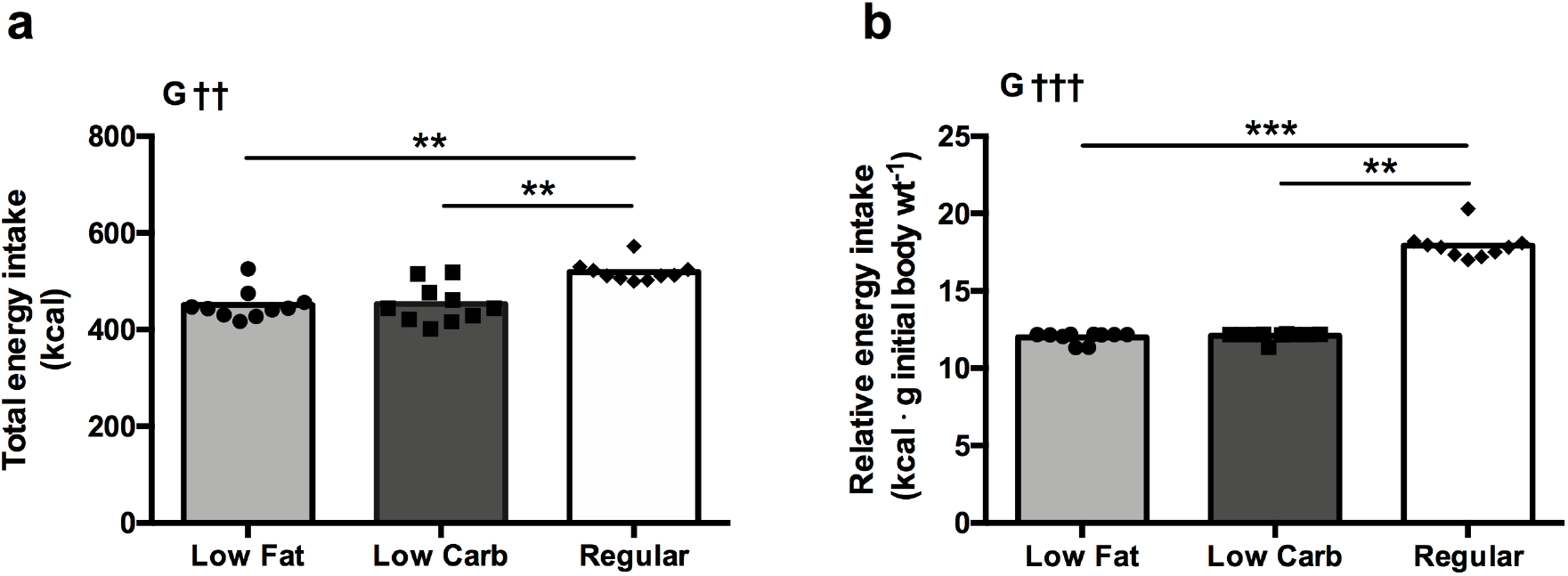
Energy intake for Low Fat and Low Carb groups during 6-week caloric restriction (CR) and in Regular group fed standard chow diet *ad libitum*. Total energy intake is shown in absolute values (a) and normalized to body mass prior to CR (b). Data are presented as mean with each dot representing one mouse data sample. Non-parametric Kruskal-Wallis with Dunn’s *post hoc* analysis was performed for group effect. †† *p* < 0.01, ††† *p* < 0.001 for group effect (G), \*\**p* < 0.01, \*\*\**p* < 0.001 between groups connected by lines.

### Body mass decreased similarly during CR in both diet groups

Data on body mass is presented in Fig. 2. Low Fat group tended to lose more weight than Low Carb group during the first week of CR (Fig. 2a). This was probably due to the reduced food intake in Low Fat group during the first week. Afterwards, however, Low Fat group caught up with food intake and showed similar weight loss as Low Carb group. Overall body mass loss did not differ between these two groups after 6-week CR (30.0 ± 5.6 and 23.8 ± 7.5% for Low Fat and Low Carb, *p* > 0.05, respectively, Fig. 2b). All mice showed clear reductions in body mass (Fig. 2c). Initially mice in the Regular diet group which were not subjected to obesogenic feeding had lower body mass (*p* < 0.001) than Low Fat and Low Carb groups, but the difference between the groups became insignificant during the final four weeks of CR which was applied to Low Fat and Low Carb groups only. Regular diet group also showed a small reduction in body mass during a final week when measurements of energy metabolism and glucose tolerance were performed after the overnight fast.

**Fig. 2.**
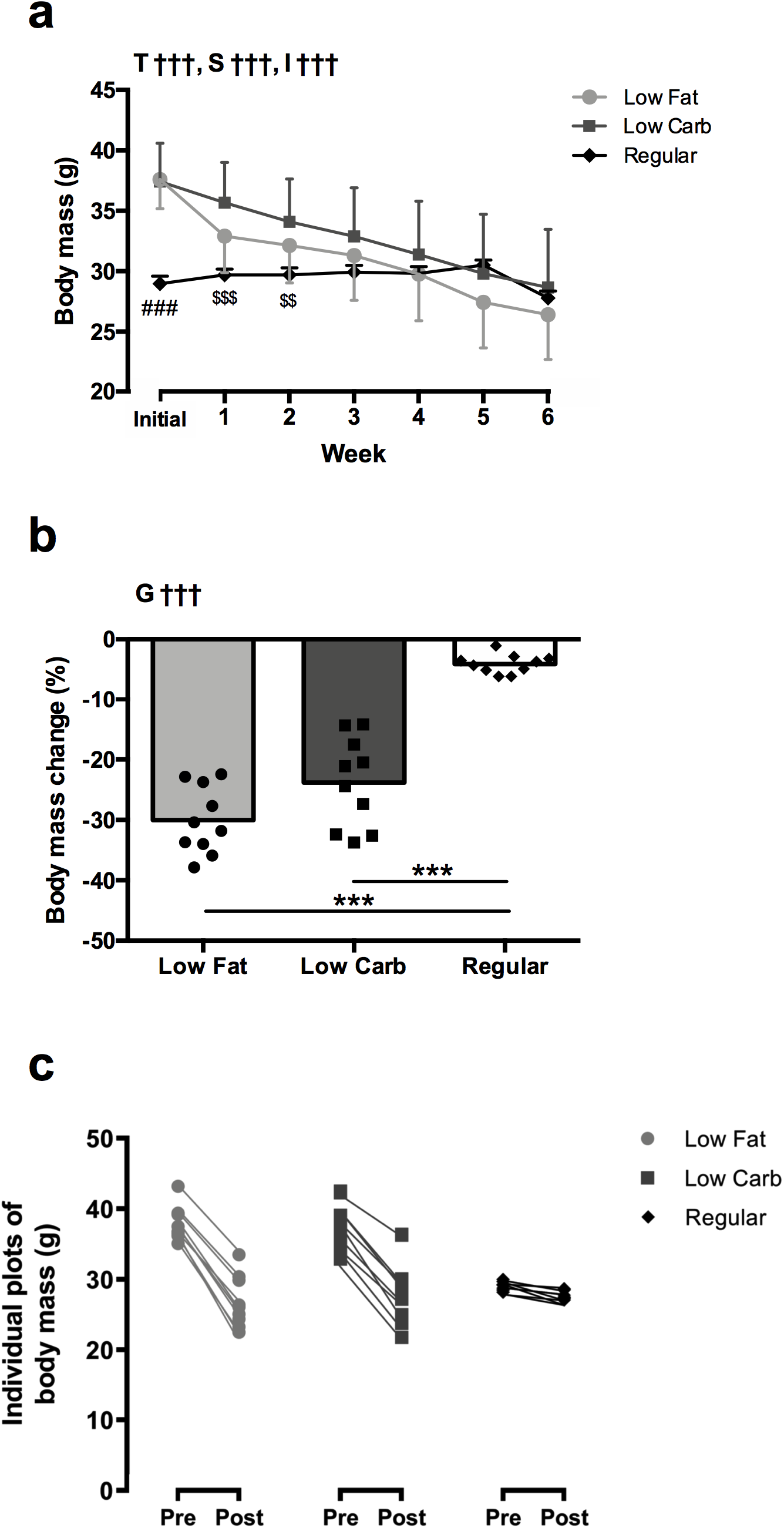
Body mass during 6-week caloric restriction (CR) in Low Fat and Low Carb diet groups as well as in Regular group fed standard chow diet *ad libitum* for the same period. Data are shown as weekly measurements (a), percentage change from initial value (b) and individual plots for the 6-week CR period (c). Data are mean ± SD (a) or mean with plotted individuals dots (b, c). Each dot represents one mouse data sample. Two-way repeated measures ANOVA (a) with Bonferroni’s *post hoc* analysis was performed for effects of group, time and subject (matching), respectively. One-way ANOVA (b) with Bonferroni’s *post hoc* analysis was performed for group effect. †† *p* < 0.01, ††† *p* < 0.001 for effects of group (G), time (T), subject (S) and interaction (I). \*\*\**p* < 0.001 between groups connected by lines, ^###^*p* < 0.001 vs. Low Fat and Low Carb, ^$$^*p* < 0.01, ^$$$^*p* < 0.001 vs. Low Carb.

### Body fat but not skeletal muscle as main energy donor during CR

Data on muscle and fat mass is shown in Fig. 3. Pre group included mice that were subjected to obesogenic diet, but did not undergo CR. This group was used to assess effects of CR on muscle and fat mass in Low Fat and Low Carb groups. Regular diet group provided age-matched reference data. Combined muscle mass differed little between the groups though it was by ~5 % smaller (*p* < 0.05) in Low Fat group compared to Pre group (Fig. 3a). Body mass normalized muscle mass increased following 6-week CR (*p* < 0.001) in Low Fat and Low Carb groups (Fig. 3b). On the other hand, body fat for these groups decreased (*p* < 0.001) to the level of Regular diet group (Fig. 3c) and became significantly lower than in Pre group (6.09 ± 2.73 and 8.57 ± 4.55 vs. 15.50 ± 3.28% body mass, *p* < 0.001, for Low Fat and Low Carb vs. Pre groups, respectively, Fig. 3d). We have also examined body fat distribution by sampling fat from five different sites of the body. Both Low Fat and Low Carb diets reduced fat mass from four out of five sites to the level of Regular diet group (Fig. 3e). An exception was iBAT which was not significantly affected by the diets and did not differ between the studied groups. Thus, CR tended to increase relative iBAT mass compared to the values prior CR (Pre group), but this increase was significant only for Low Fat group (*p* < 0.01) (Fig. 3f). Relative mass of gWAT decreased more in Low Fat than Low Carb group (*p* < 0.05).

**Fig. 3.**
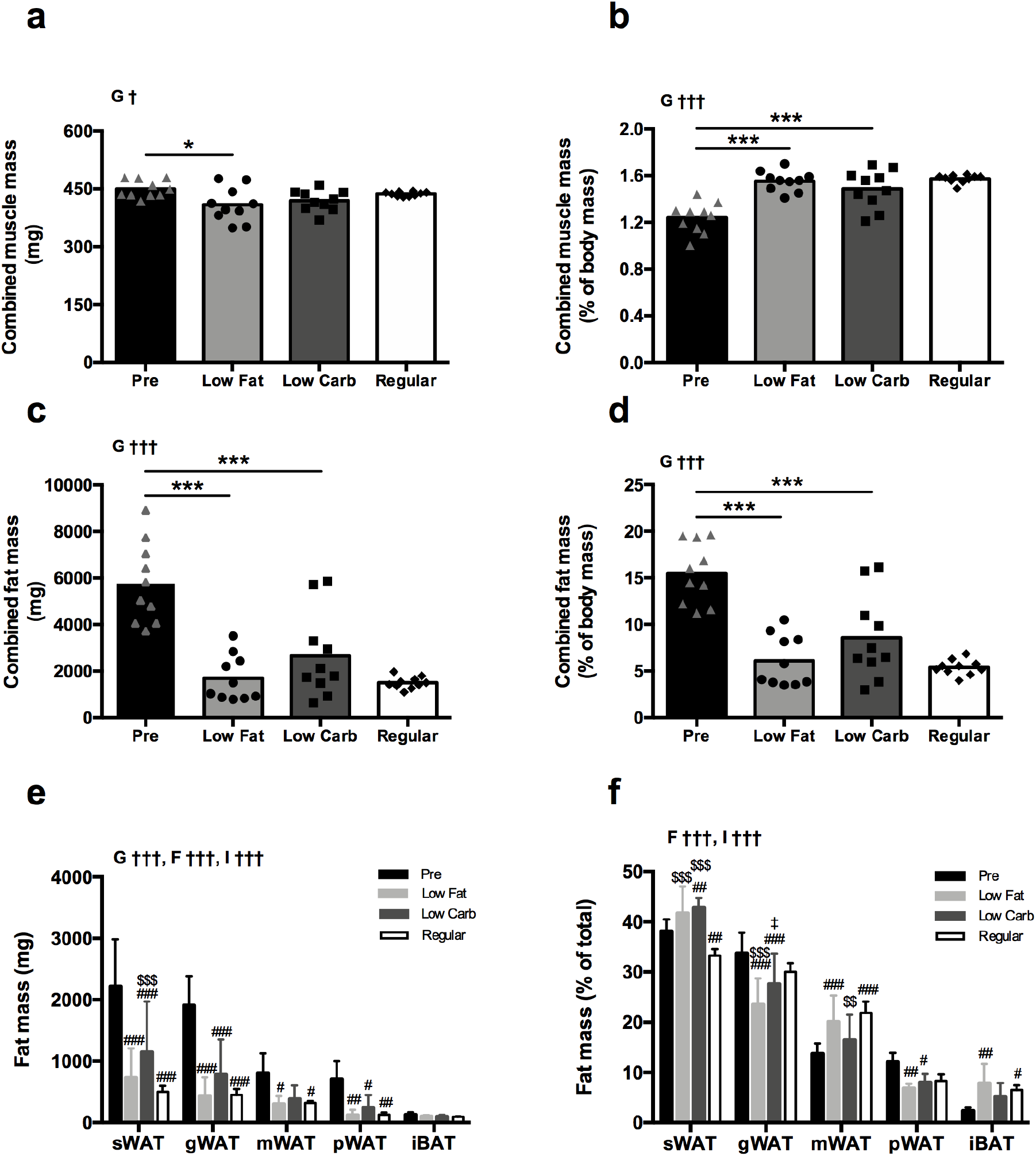
Mass changes of skeletal muscle (a, b), body fat (c, d) and fat from different sampling sites (e, f) in Low Fat and Low Carb diet groups after 6-week caloric restriction (CR) compared to the obese group prior CR (Pre) as well as to the age-matched Regular group fed standard chow diet *ad libitum* for the same period. Abbreviations (e, f): sWAT, subcutaneous white adipose tissue; gWAT, gonadal white adipose tissue; mWAT, mesenteric white adipose tissue; pWAT, perirenal white adipose tissue; iBAT intrascapular brown adipose tissue. Data are mean ± SD (e, f) or mean with plotted individuals dots (a, b, c, d). Each dot represents one mouse data sample. One-way ANOVA (a-d) with Bonferroni’s *post hoc* analysis was performed for group effect. Two-way repeated measures ANOVA (e, f) with Bonferroni’s *post hoc* analysis was performed for effects of group and fat site, respectively. † *p* < 0.05, ††† *p* < 0.001 for effects of group (G), fat site (F) and interaction (I). \**p* < 0.05 and \*\*\**p* < 0.001 between groups connected by lines, ^#^*p* < 0.05, ^##^*p* < 0.01 and ^###^*p* < 0.001 vs. Pre, ^$$^*p* < 0.01, ^$$$^*p* < 0.001 vs. Regular; ^‡^*p* < 0.05 vs. Low Fat.

### Glucose tolerance improves similarly independently of the diets after CR

Data on glucose tolerance from IPGTT is presented in Fig. 4. Glucose AUC was similar in Low Fat and Low Carb groups (*p* > 0.05), but smaller compared to Pre group (*p* < 0.01) and Regular diet group (*p* < 0.05) (Fig. 4b). Pre and Regular diet groups did not differ in glucose AUC though Regular diet group demonstrated a large initial spike with subsequent normalization of blood glucose to baseline values whereas Pre group showed slow rise in blood glucose which did not show any decrease during the entire 2 h duration of the test (Fig. 4a).

**Fig. 4.**
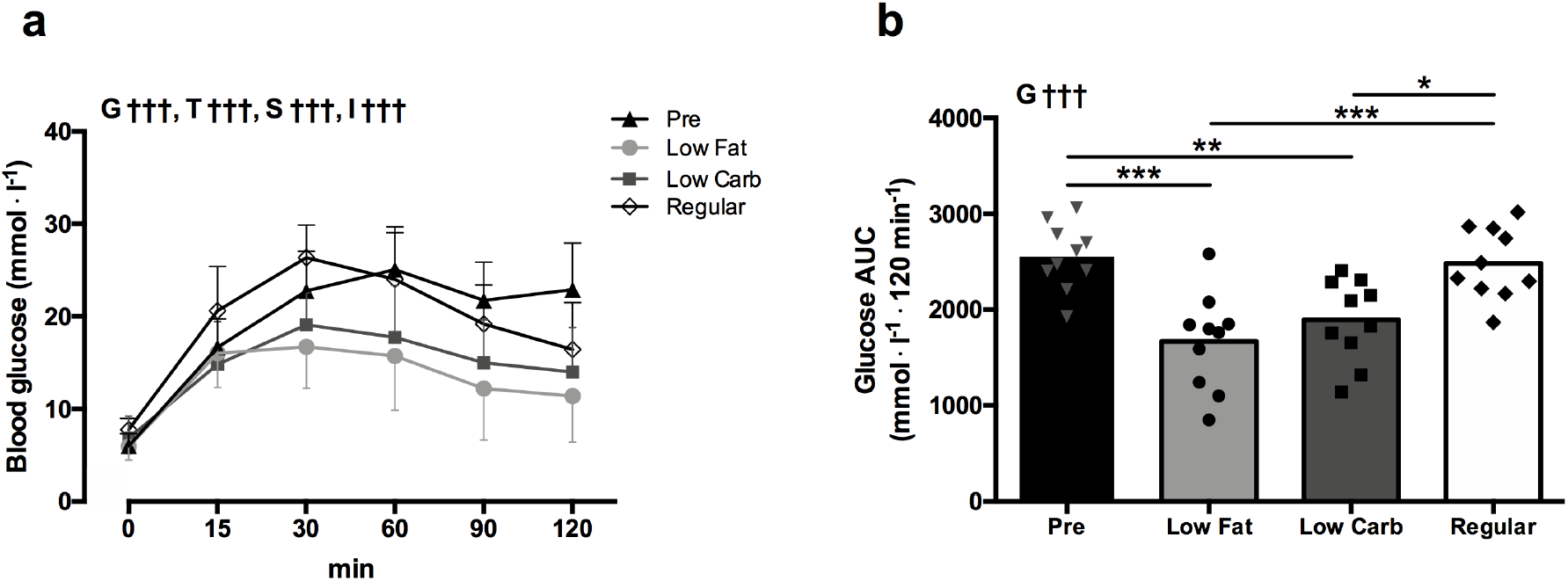
Pattern of blood glucose clearance during a 120 min period (a) and glucose area under curve (AUC) (b) in Low Fat and Low Carb diet groups after 6-week caloric restriction (CR) compared to the obese group prior CR (Pre) as well as to the age-matched Regular group fed standard chow diet *ad libitum* for the same period. Data are mean ± SD (a) or mean with plotted individuals dots (b). Each dot represents one mouse data sample. Two-way repeated measures ANOVA (a) with Bonferroni’s *post hoc* analysis was performed for effects of group, time and subject (matching), respectively. One-way ANOVA (b) with Bonferroni’s *post hoc* analysis was performed for group effect. ††† *p* < 0.001 for effects of group (G), time (F), subject (S) and interaction (I). \**p* < 0.05, \*\**p* < 0.01 and \*\*\**p* < 0.001 between groups connected by lines.

### Low Fat and Low Carb diets had the same effect on energy metabolism and physical activity

Data on energy metabolism are presented in Fig. 5. Total energy expenditure did not differ between Low Fat and Low Carb groups (*p* < 0.05) (Fig. 5a). Pre group showed higher (*p* < 0.05) energy expenditure than Regular diet group, but ANCOVA analysis with body mass and physical activity as covariates did not show any significant differences between the groups and showed that physical activity but not body mass had an effect on energy expenditure. There were no significant differences in physical activity between the groups which was probably due to rather large variations within the groups. Low Fat and Low Carb groups tended to be more active than Pre or Regular diet groups (Fig. 5b). Association between physical activity and energy expenditure was significant in all groups (r = 0.70-0.80, *p* < 0.05-0.01) (Fig. 5c). Linear regression analysis showed a tendency for a slightly greater predictive resting metabolic rate in Low Carb compared to Low Fat group (0.35 vs. 0.30 kcal · h^−1^). On the other hand, respiratory quotient did not differ between the groups when measurements were performed in the fasted mice (Fig. 5d).

**Fig. 5.**
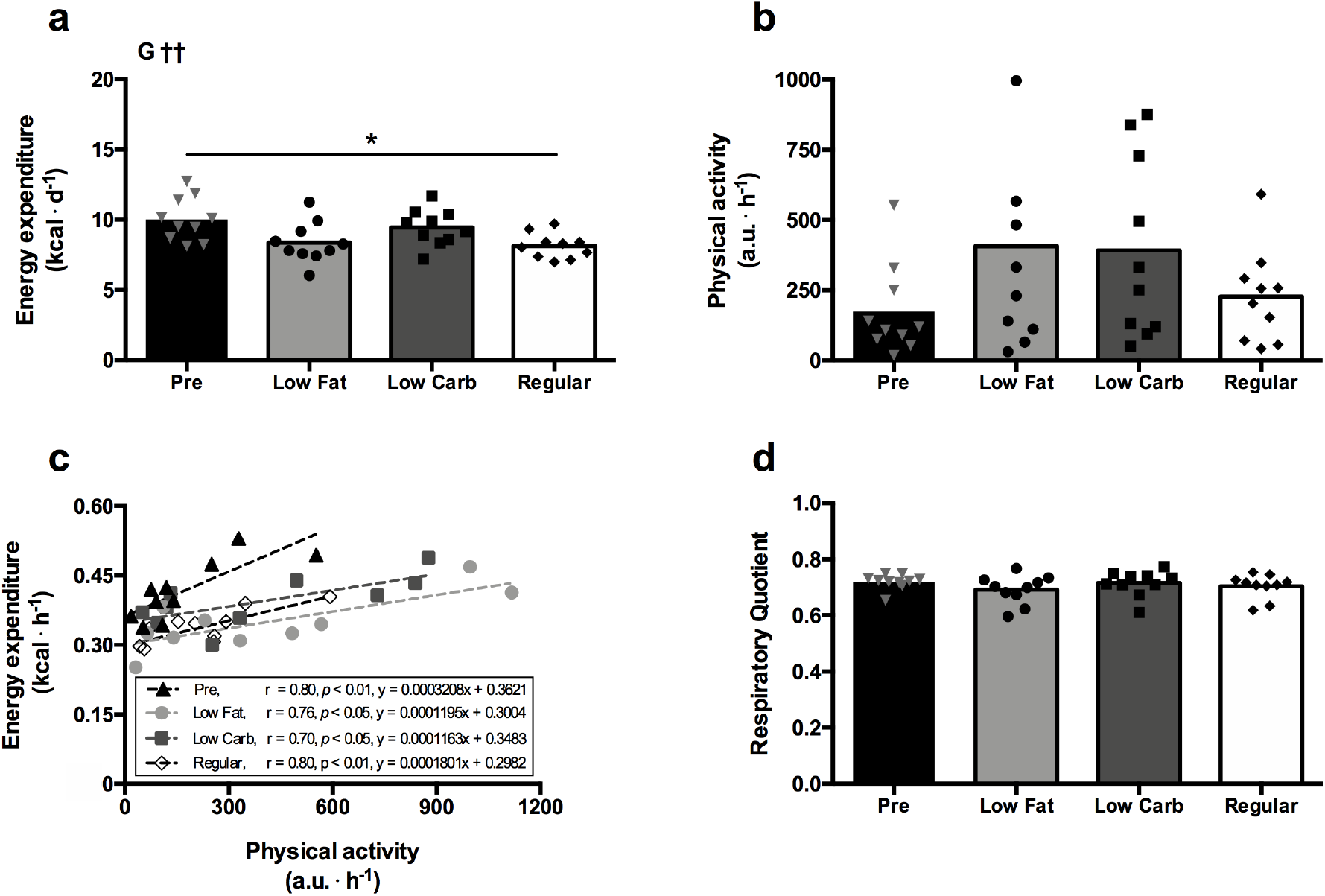
Total energy expenditure, (a), physical activity (b), plots of energy expenditure versus physical activity (c) and respiratory quotient (d) in Low Fat and Low Carb diet groups after 6-week caloric restriction (CR) compared to the obese group prior CR (Pre) as well as to the age-matched Regular group fed standard chow diet *ad libitum* for the same period. All the measurements were performed after overnight fasting. Data are presented as mean with each dot representing one mouse data sample. One-way ANOVA (a, b, d) with Bonferroni’s *post hoc* analysis was performed for group effect. †† *p* < 0.01 for group effect (G), \**p* < 0.05 between groups connected by lines. Pearson correlation coefficient (r) and linear regression equations for the plots of energy expenditure versus physical activity are also shown (c).

## Discussion

The main aim of our study was to investigate if carbohydrate and fat content of diets affects physiological responses to caloric restriction in mice. Most of the previous studies have focused on effects of macronutrient content of diets on health and body composition in *ad libitum* fed mice (23, 31) and there are only few studies under conditions of caloric restriction (32). In agreement with recent findings on humans undergoing mild caloric restriction (33), our results show that improvements in body composition and glucose tolerance do not differ between low fat and low carbohydrate diets with equal protein content under conditions of up to 40% caloric restriction. This is important in view of the fact that differences between low carbohydrate and low fat diets have been widely discussed in relation to health outcomes (34).

Carbohydrate-insulin model of obesity has been proposed in justification of health benefits of low carbohydrate diets (14). According to this line of reasoning high carbohydrate content of food leads to high blood insulin levels which acts to suppress the release of fatty acids from adipose tissue and directs circulating fat towards adipose tissue for storage rather than oxidation in metabolically active tissues. However, a large number of studies contradicts the carbohydrate-insulin model of obesity. Meta-analysis of 32 controlled feeding studies with substitution of carbohydrate for fat show that fat loss and energy expenditure were greater for low fat diets compared to low carbohydrate diets though differences between the diets in fat loss (16 g · d^−1^) and energy expenditure (26 kcal · d^−1^) were rather small (35). A recent randomized clinical trial which engaged over 600 participants showed no difference between low fat and low carbohydrate diets in weight loss during a 12-month period, and neither baseline insulin secretion nor genotype pattern was associated with the dietary effects on weight change (33). All the above-mentioned studies controlled energy intake and equated dietary protein between diets. It appears that protein is a macronutrient which is particularly important for dietary-induced thermogenesis and satiety. The thermic effect of dietary protein is 25-30% of its energy content compared to 5-10% and 2-3% for carbohydrate and fat, respectively (12). Increase in protein intake from 15 to 30% of total energy is associated with spontaneous reduction in total energy intake under conditions of *ad libitum* feeding (36). High protein intake also led to increase in retention of lean body mass during caloric restriction (37). Thus, comparison of high fat and high carbohydrate diets can be compromised by differences in protein content, as health benefits of protein-rich diets are often incorrectly assigned to carbohydrate and fat content of the diets (38). In our study, we kept both the amount (20% of total energy intake) and source (casein with addition of l-cystine) of dietary protein constant between the low fat and low carbohydrate diets. It appears that this amount of protein was adequate for skeletal muscle mass retention which did not change significantly during caloric restriction. Furthermore, mice were fed obesogenic diet for 18-week prior to caloric restriction. This diet induces minor changes in lean body and significant increase in body fat which might also help to preserve muscle mass during caloric restriction (25). Human weight loss studies show that approximately 25% of weight loss is due to loss of lean body mass with major contribution of the skeletal muscles to this decline (39). People who are leaner tend to lose more of lean body mass under conditions of caloric restriction compared to those with greater body fat content (40). It appears that obese mice show greater sparing of muscle mass during caloric restriction compared to humans. However, dissection of factor playing a role in preservation of muscle mass in mice and/or humans during caloric restriction was beyond the scope of our current study.

It appears that body fat was the main source of energy during caloric restriction and its loss did not differ between the two diets in our study. Increased fatty acid oxidation is a common feature of low carbohydrate high fat diets which are often perceived as more lipolytic and less obesogenic compared to low fat high carbohydrate diets though human metabolic ward studies challenges this hypothesis (41). We did not observe any differences between the diets in respiratory quotient as the measurements were performed in the fasted state. Mice gorge on food and consume all the food within 2-4 h period of time after feeding when subjected to caloric restriction (42, 43). Thus, when exposed to caloric restriction, mice spent significant periods of time in the fasted state which is associated with high rate of fatty acid oxidation (28).

Thus, measurements in the fasted state might be more representative of the overall metabolism compared to measurements in the post-absorptive state under conditions of caloric restriction. It appears that metabolic flexibility manifesting itself in switching between carbohydrate and fat oxidation allowed to maintain a similar net body fat balance in mice independently of the macronutrient composition of the diets during caloric restriction (44).

Linear regression analysis of the plots for physical activity over energy expenditure allowed to exclude effects of physical activity on energy expenditure and showed that predicted resting metabolic rate tended to be slightly greater under conditions of low carbohydrate diet compared to the low fat diet. However, this difference between the diets was not significant and can hardly be used as evidence in support of recent findings in human studies that low carbohydrate diets lead to greater energy expenditure compared to low fat diets (16). Our results are in agreement with many human studies that reported no practically meaningful differences in energy expenditure between the isocaloric and isonitrogenous low fat and low carbohydrate diets (33, 35, 45).

We have assessed glucose tolerance as a key indicator of metabolic health (46). After 6-week caloric restriction glucose tolerance improved similarly in both diets. It is likely that caloric restriction-induced loss of body fat was a key factor promoting better glucose control irrespective of dietary carbohydrate and fat content. In contrast to our findings, a recent caloric restriction study of C57BL/6 mice showed smaller improvement in glucose tolerance for high fat diet compared to chow diet which is high in carbohydrates in spite of similar weight loss for both diets (32). However, macronutrients and their sources were not strictly controlled in the latter study and the protein content differed substantially between both diets, i.e. 20% kcal for high fat diet and 33% kcal for chow diet low in fat. Dietary protein due to its insulinotropic effects might potentially influence postprandial glucose control (47, 48). There is evidence that consumption of the high protein meal before the intake of carbohydrates attenuates the subsequent rise in the postprandial serum glucose and results in lower glucose compared to isocaloric high carbohydrate and high fat meals (49). In humans, weight loss is a priority target under the conditions of impaired glucose homeostasis as in case of type 2 diabetes (50). Antidiabetic therapies that can control blood glucose levels but promote weight gain are less effective as greatest improvements in glucose control are observed in patients with greatest reductions in body mass (51). Taken together, caloric restriction-induced body fat loss should be considered as a primary and most desirable target for positive management of blood glucose whereas macronutrient composition of isocaloric diets with equated protein probably plays a minor role at its best.

In summary, isocaloric energy restriction with equated dietary protein rather than a distribution of dietary carbohydrate and fat was a main factor for favourable changes of body composition in obese mice. Improvements of blood glucose control in obese mice was driven by body fat loss irrelevant to dietary carbohydrate and fat ratio in the diet. It appears that the overall energy and dietary protein intake should be targeted when the aim is to improve body composition and glucose control while dietary carbohydrate and fat content should be left to personal preference for adherence purposes.

## Acknowledgments

We would like to thank Mrs Indrė Libnickienė for excellent technical assistance during the project.

## Author Contributions

P.M. conceived and designed research; P.M., A.F., M.D. performed experiments; P.M. A.F. and A.R. analysed data; P.M., A.F. and A.R. interpreted results of experiments; P.M. and A.F. prepared figures; P.M. and A.F. drafted manuscript; P.M., A.F. and A.R. edited and revised manuscript; P.M., A.F., M.D. and A.R. approved final version of manuscript.

---

**Supplementary table 1.** Detailed macronutrient composition of the diets provided by a manufacturer (Research Diets Inc.)

| Diet # | Obesogenic diet<br>(D12451, Research<br>Diets Inc., USA) |  | Low Fat<br>(D17100401, Research<br>Diets Inc., USA) |  | Low Carbohydrate<br>(D12492, Research<br>Diets Inc., USA) |  |
| --- | --- | --- | --- | --- | --- | --- |
|  | g (%) | kcal (%) | g (%) | kcal (%) | g (%) | kcal (%) |
| Protein | 24 | 20 | 20 | 20 | 26 | 20 |
| Carbohydrate | 41 | 35 | 61 | 60 | 26 | 20 |
| Fat | 24 | 45 | 9 | 20 | 35 | 60 |
| Total |  | 100 |  | 100 |  | 100 |
| kcal · g<sup>-1</sup> | 4.73 |  | 4.1 |  | 5.2 |  |
| **Ingredients** |  |  |  |  |  |  |
| **Protein:** |  |  |  |  |  |  |
| Casein, 30 Mesh | 200 | 800 | 200 | 800 | 200 | 800 |
| L-Cystine | 3 | 12 | 3 | 12 | 3 | 12 |
| **Carbohydrate:** |  |  |  |  |  |  |
| Corn Starch | 72.8 | 291 | 405 | 1620 | 0 | 0 |
| Maltodextrin 10 | 100 | 400 | 125 | 500 | 125 | 500 |
| Sucrose | 172.8 | 691 | 68.8 | 275 | 68.8 | 275 |
| **Fibre:** |  |  |  |  |  |  |
| Cellulose, BW 200 | 50 | 0 | 50 | 0 | 50 | 0 |
| **Fat:** |  |  |  |  |  |  |
| Soybean Oil | 25 | 225 | 25 | 225 | 25 | 225 |
| Lard | 177.5 | 1598 | 65 | 585 | 245 | 2205 |
| **Minerals:** |  |  |  |  |  |  |
| Mineral Mix S10026 | 10 | 0 | 10 | 0 | 10 | 0 |
| DiCalcium Phosphate | 13 | 0 | 13 | 0 | 13 | 0 |
| Calcium Carbonate | 5.5 | 0 | 5.5 | 0 | 5.5 | 0 |
| Potassium Citrate, 1<br>H<sub>2</sub>O | 16.5 | 0 | 16.5 | 0 | 16.5 | 0 |
| **Vitamins:** |  |  |  |  |  |  |
| Vitamin Mix V10001 | 10 | 40 | 10 | 40 | 10 | 40 |
| Choline Bitartrate | 2 | 0 | 2 | 0 | 2 | 0 |
| **Total** | **858.15** | **4057** | **998.75** | **4057** | **773.85** | **4057** |

